# Glycolytic metabolism modulation on spinal neuroinflammation and vital functions following cervical spinal cord injury

**DOI:** 10.1101/2024.05.14.594105

**Authors:** Pauline Michel-Flutot, Arnaud Mansart, Stéphane Vinit

## Abstract

High spinal cord injuries (SCIs) often result in persistent diaphragm paralysis and respiratory dysfunction. Chronic neuroinflammation within the damaged spinal cord after injury plays a prominent role in limiting functional recovery by impeding neuroplasticity. In this study, we aimed to reduce glucose metabolism that supports neuroinflammatory processes in an acute preclinical model of C2 spinal cord lateral hemisection in rats. We administered 2-deoxy-D-glucose (2-DG; 200 mg/kg/day s.c., for 7 days) and evaluated the effect on respiratory function and chondroitin sulfate proteoglycans (CSPGs) production around spinal phrenic motoneurons. Contrary to our initial hypothesis, our 2-DG treatment did not have any effect on diaphragm activity and CSPGs production in injured rats, although slight increases in tidal volume were observed. Unexpectedly, it led to deleterious effects in uninjured (sham) animals, characterized by increased ventilation and CSPGs production. Ultimately, our results seem to indicate that this 2-DG treatment paradigm may create a neuroinflammatory state in healthy animals, without affecting the already established spinal inflammation in injured rats. Given the beneficial effects of 2-DG observed in other studies on neuronal activity and inflammation, adjusting 2-DG doses and/or increasing treatment duration should be explored to reduce deleterious inflammatory processes occurring after SCI.

## Introduction

Spinal cord injuries (SCI) occurring at high cervical levels result in a range of functional impairments, including persistent respiratory disorders (Winslow and Rozovsky, 2003; Ahuja et al., 2017). Cervical SCIs are the most prevalent, and patients living with these injuries often have to rely on mechanical ventilation to survive, and are thus more susceptible to respiratory infections (Ahuja et al., 2017). To date, no effective treatments that induce complete functional recovery are available, making the development of new therapeutics critical (Randelman et al., 2021; Locke et al., 2022; Michel-Flutot et al., 2023a).

One of the most commonly used preclinical models to study respiratory function following high SCI is the C2 spinal cord hemisection model (C2HS). This model also allow to study SCI impact on spinal neuroplasticity and neuroinflammation related to the phrenic system (Porter, 1895; Nantwi et al., 1999; Golder et al., 2001a; Vinit et al., 2006; Lane et al., 2009; Keomani et al., 2014). A C2HS leads to a disruption of half of the descending respiratory fibers ipsilateral to the injury. These fibers originate from the rostral ventral respiratory group in the brainstem and project to the phrenic motoneurons, located in spinal cord segments C3-C6. Their axons form the phrenic nerve which innervates the diaphragm, the primary inspiratory muscle. This injury results in diaphragm hemiplegia ipsilateral to the initial injury, while the contralateral side remains intact and allows the animal to survive (Vinit et al., 2006; Vinit and Kastner, 2009; Keomani et al., 2014; Lee et al., 2014; Allen et al., 2021; Rana et al., 2022).

In this C2HS preclinical model, spontaneous (though limited) recovery of diaphragm activity can be observed. This is due to the crossed phrenic phenomenon (CPP), characterized by a partial reactivation of the phrenic network in the C3-C6 spinal cord sustained by previously silent pathways crossing the midline at the level of phrenic motoneurons (Fuller et al., 2006; Goshgarian, 2009; Ghali, 2017). However, this marginal plasticity does not account for any significant ventilatory recovery (Dougherty et al., 2012).

In addition to the functional impairments they cause, SCIs also include pathological inflammatory components in a time-dependent manner (acute to chronic phase) (Fawcett et al., 2012; Cregg et al., 2014; Bradbury and Burnside, 2019). For instance, a variety of molecules are expressed or overexpressed in the context of chronic inflammation, which in turn contributes to the inhibition of neuronal plasticity observed following SCI (Bradbury and Burnside, 2019). Chondroitin sulfate proteoglycans (CSPGs) are molecules present in the perineuronal network (PNN) (Bartus et al., 2012; Kwok et al., 2014; Bradbury and Burnside, 2019). CSPGs can bind different receptors involved in neuronal growth inhibition (Bradbury and Burnside, 2019), such as PTPσ receptor (Shen et al., 2009), leukocyte common antigen-related phosphatase (LAR) (Fisher et al., 2011), and Nogo receptors (NgR1 and NgR3) (Dickendesher et al., 2012). The interaction between CSPGs and their receptors activates intracellular signaling pathways that inhibit axonal growth, preventing axonal plasticity and functional recovery following SCI (Bradbury and Burnside, 2019). These CSPGs are mainly overexpressed by reactive astrocytes after SCI (Bradbury and Burnside, 2019), therefore CSPGs expression can therefore be related to how permissive the PNN will be to axonal growth, as well as to the level of inflammation. Furthermore, several works demonstrated that removal of CSPGs lead to improvements in motor function (Bradbury et al., 2002; Tester and Howland, 2008; Kwok et al., 2014; James et al., 2015; Warren et al., 2018; Muir et al., 2019; Rosenzweig et al., 2019). Therefore, reducing CSPG expression surrounding phrenic motoneurons in the injured spinal cord after C2HS may result in similar improvements in respiratory function.

The molecule 2-deoxy-D-glucose (2-DG) is an exogen glucose mimic which can be used to competitively and partially inhibit glycolysis (Woodward and Hudson, 1954; Xi et al., 2014). 2-DG is molecularly identical to glucose, except the hydroxyl group present in position 2 of glucose is replaced by a hydrogen in 2-DG. Like glucose, 2-DG enters cells via glucose transporter proteins (GLUTs), where it competes with glucose for binding to hexokinase.

Hexokinase converts 2-DG into 2-deoxy-D-glucose-6-phosphate, which accumulates in the cell as it cannot be further metabolized. This leads to a reduction in glucose metabolism in dose-dependent fashion, and consequently, a reduction of glycolysis and pentose phosphate products (Xi et al., 2014; Dey et al., 2022).

2-DG is mainly studied for its potential as an anticancer agent due to its inhibitory effects on glycolysis. This results in nutrient and energy deprivation in cancer cells, leading to a subsequent reduction in growth and survival (Pajak et al., 2020; Dey et al., 2022). 2-DG has also been instrumental in elucidating the role glucose metabolism plays in the context of various neuropathological states. For instance, a reduction in cellular glucose intake caused by 2-DG leads to a delay and reduction in the frequency of seizures *ex vivo* and *in vivo*, making 2-DG a potential therapeutic treatment for epileptic episodes (Stafstrom et al., 2008; Sutula et al., 2023). 2-DG also has broad anti-inflammatory properties (Dey et al., 2022). For instance, glucose is involved in macrophage polarization (Obaid et al., 2021), and the formation of hypoxia-inducible factor 1-alpha (HIF-α), which is involved in pro-inflammatory macrophage signaling (Ip et al., 2017; Dey et al., 2022). Therefore, by inhibiting glycolysis, 2-DG administration can reduce these inflammatory phenomena (Dey et al., 2022). 2-DG treatment has also been shown to lead to a reduction of sevoflurane induced-neuroinflammation *in vitro*, specifically in mouse primary microglia cells. This treatment lead to reductions in the levels of Interleukin (IL)-6, tumor necrosis factor (TNF-α) and nuclear factor-kappa B (NFκB) (Wang et al., 2014). Similarly, treatment of LPS induced-inflammation in microglia with 2-DG induces a decrease in HIF-1α accumulation and IL-1β production, attributable to the inhibition of glycolysis (York et al., 2021).

Given the beneficial effects of 2-DG on inflammatory processes, and the involvement of neuroinflammation in neuroplasticity inhibition following SCI, we hypothesized that administration of 2-DG in a preclinical model of C2HS would lead to beneficial effect on ventilatory function, as well as on neuroinflammatory processes, characterized by CSPGs expression deposition in the spinal cord.

## Material and Methods

### Ethics statement

All experiments reported in this manuscript conformed to policies laid out by the National Institutes of Health (Bethesda, MD, USA) in the Guide for the Care and Use of Laboratory Animals and the European Communities Council Directive of 22 September 2010 (2010/63/EU, 74) regarding animal experimentation (Apafis #2017111516297308_v3). These experiments were performed on 3–4-month-old male Sprague–Dawley rats (Janvier, France). The animals were dual-housed in individually ventilated cages in a state-of-the-art animal care facility (2CARE animal facility, accreditation A78-322-3, France), with access to food and water *ad libitum* with a 12 h light/dark cycle.

### Chronic C2 hemisection and 2-deoxy-D-glucose administration

In this study, 19 animals were used, divided into 4 groups: Sham animals treated with saline (*n* = 3) or 2-DG (*n* = 4) for 7 days post-laminectomy (sham surgery), and animals treated with saline (*n* = 8) or 2-DG (*n* = 4) for 7 days post-injury (P.I.). All animals were anesthetized with isoflurane (Iso-vet, Piramal, Voorschoten, Netherlands; ∼1.5% in O_2_) before receiving bilateral 15 μL intrapleural injections of the retrograde tracer cholera toxin B fragment at a concentration of 0.2% (CTB, List Biologicals, Campbell, CA, USA) to identify phrenic motor neurons in the cervical spinal cord as previously described (Mantilla et al., 2009; Michel-Flutot et al., 2022). Three days after CTB injection, rats underwent C2HS or sham surgery. As described previously (Keomani et al., 2014), buprenorphine (Buprécare, 0.03 mg/kg), trimethoprim, and sulfadoxine (Borgal 24%, 30 mg/kg), medetomidine (Médétor, 0.1 mg/kg) and carprofen (Rimadyl, 5 mg/kg) were administered subcutaneously 10–20 min before inducing isoflurane anesthesia in a closed chamber (5% isoflurane in 100% O_2_). Rats were orotracheally intubated and ventilated with a rodent ventilator (model 683; Harvard Apparatus, South Natick, MA, USA). Anesthesia was maintained throughout the procedure (1.5–2% isoflurane in 100% O_2_). Skin and muscles were retracted, and a C2 laminectomy and durotomy were performed. For the C2-injured animals, the spinal cord was hemisected unilaterally just caudal to the C2 dorsal roots with micro-scissors, followed with a micro-scalpel to ensure the section of all the remaining fibers, as described previously by our team (Keomani et al., 2014). The wounds and skin were then sutured. Sham rats underwent the same procedures without spinal cord hemisection. At the conclusion of the procedures, an intra-muscular injection of atipamezole (0.5 mg/kg, Revertor, Virbac, Carros, France) was administered to reverse medetomidine. The isoflurane was discontinued, the endotracheal tube was pulled off, and the rats were monitored throughout recovery. Rats were subcutaneously injected with 2-DG (200 mg/kg/day, Carbosynth, MD05187-1101) or saline (NaCl 0.9%) every 24-hours for 7 consecutive days starting on the day of surgery.

### Whole-body plethysmography

Whole-body constant flow (2 L/min) plethysmography (EMKA, France) was used at 7-days post-surgery to assess global ventilatory function (e.g., the tidal volume, minute ventilation, and breathing frequency) in our rats, as previously described (Fayssoil et al., 2021). The rats were weighed, and the plethysmography system was calibrated before the animals were placed in the chambers. After a 30-minute acclimation period, recording began under normoxic conditions (room air). The minute ventilation and tidal volume were reported according to bodyweight (per 100 g) for each animal.

### Electrophysiological and hemodynamic recordings

All animals were then assessed for their diaphragm activity. Briefly, anesthesia was induced using isoflurane (5% in 100% air balanced) in an anesthesia chamber and maintained through a nose cone (2.5% in 100% air balanced). Animals were placed on a heating pad to maintain a constant body temperature (37.5 ± 0.5 °C), and their rectal temperature was continuously monitored throughout the experiment. The depth of anesthesia was confirmed by the absence of response to toe pinch. Arterial pressure and heart rate were measured through a catheter inserted into the right femoral artery. Arterial pressure was monitored continuously with a transducer connected to a bridge amplifier (AD Instruments, Dunedin, New Zealand). A laparotomy was performed, and the liver was gently moved dorsally to access the diaphragm. Gauze soaked with warm phosphate-buffered saline was placed on the liver to prevent dehydration. A handmade bipolar surface silver electrode was used to record spontaneous diaphragm EMG during poïkilocapnic normoxic or transient mild asphyxia breathing (by occlusion of the animal’s nose for 15 s, as published before (Michel-Flutot et al., 2022)).

EMGs were amplified (Model 1800; gain, 100; A-M Systems, Everett, WA, USA) and band pass-filtered (100 Hz to 10 kHz). The signals were then digitized with an 8-channel Powerlab data acquisition device (Acquisition rate: 4 k/s; AD Instruments, Dunedin, New Zealand) connected to a computer and analyzed using LabChart 8 Pro software (AD Instruments, Dunedin, New Zealand). The bilateral diaphragmatic EMGs were twice integrated (50 ms constant decay) in order to reduce the parasitic electrocardiogram signal.

### Tissue Processing

At the end of the experiment, animals were euthanized by intracardiac injection of pentobarbital (EXAGON, Axience), intracardially perfused with heparinized 0.9% NaCl followed by Antigenfix solution (DIAPATH, Martinengo (BG), Italy). After perfusion, the C1-C6 spinal cord was carefully dissected and stored at 4°C in fixative for 24h. After post-fixation, tissues were cryoprotected for 48h in 30% sucrose (in 0.9% NaCl) and stored at -80°C. Frozen longitudinal (C1-C3 spinal cord) and transverse (C3-C6 spinal cord) free floating sections (30 µm) were cut using a cryostat (Thermo Fisher). C3-C6 sections were stored in a cryoprotectant solution (Sucrose 30%, ethylene glycol 30% and PVP40 1% in PBS 1X) at −22°C. Every fifth section from C1-C3 were used for lesion reconstruction to examine the extent of C2 injury using cresyl violet histochemistry.

### Histological reconstruction of the extent of C2 injury

Longitudinal sections from C1-C3 cord were used to assess the dorso-ventral and medio-lateral extent of injury in all animals. Brightfield microscopy was used to examine the cresyl violet stained sections and the lesion area was recorded on a stereotaxic transverse plane of the C2 spinal cord. Each injury was then digitized and analyzed with Image J software (NIH). The extent of the injury on the injured side was calculated using a reference to a complete hemisection (which is 100% of the hemicord) and reported as a percentage of the hemicord as we published before (Keomani et al., 2014; Michel-Flutot et al., 2022) (Figure 1).

**Figure 1.**
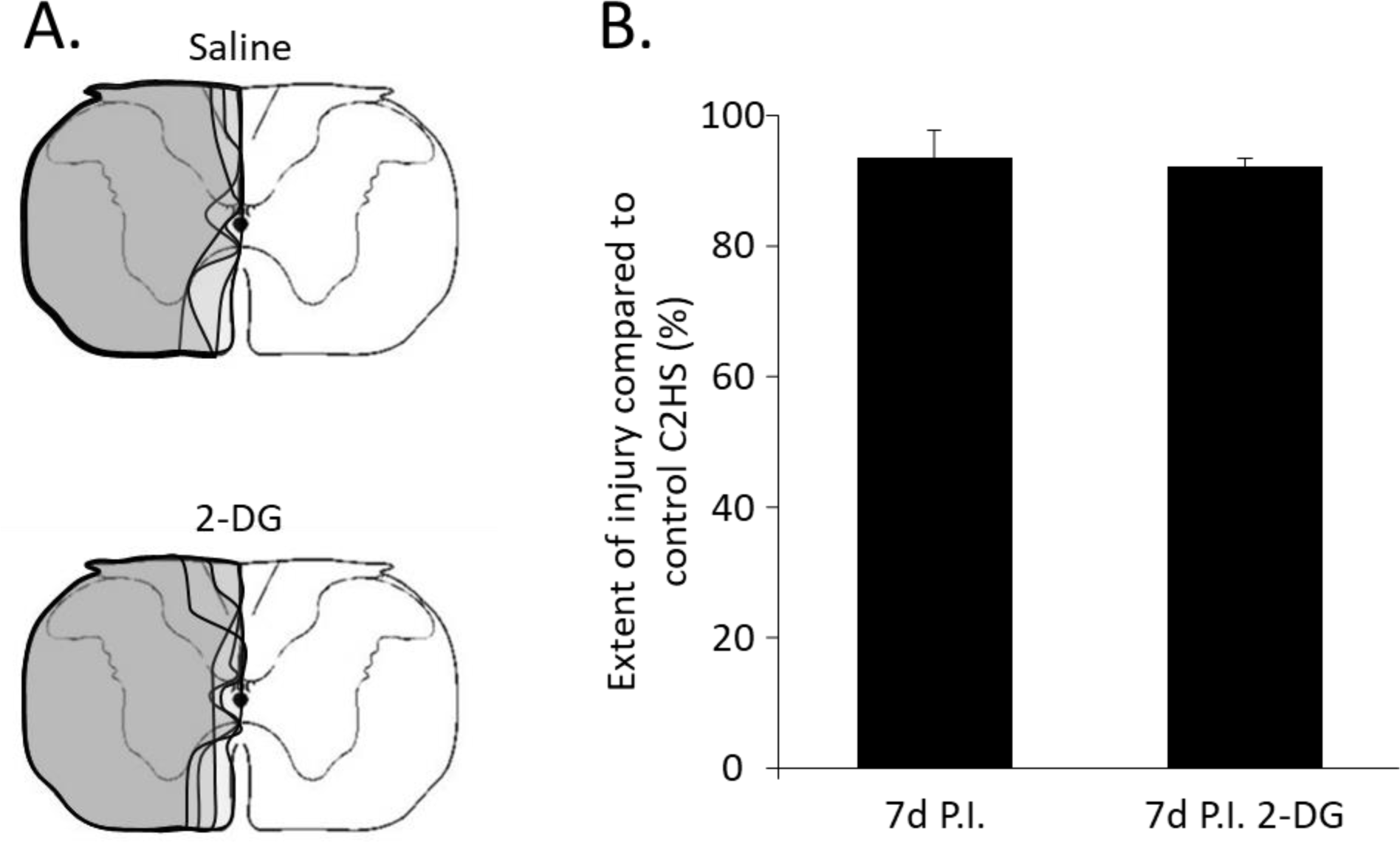
Extent of injury following a C2 spinal cord hemisection. (A) Representative extent of injury in each animal at 7 days post-injury (P.I.) for saline and 2-deoxy-D-glucose (2-DG) treated groups; (B) Extent of injury quantification compared to a full spinal cord C2 hemisection (C2HS; 100%) in percentage. The quantification was done only for the ventral part of the spinal cord where descending respiratory pathways are located (Vinit et al., 2006). No difference was observed between groups (Student’s t-test, p = 0.591).

### Immunofluorescence

For immunofluorescence analyses, 13 animals (out of the 19 used in this study) were used: 7-day post-surgery sham animals treated with saline (*n* = 2) or 2-DG (*n* = 3), and 7-day post-injury (P.I.) animals treated with saline (*n* = 4) or 2-DG (*n* = 4). Free-floating transverse sections of the C3-C6 spinal cord were washed and placed in blocking solution (Normal Donkey Serum (NDS) 5% and Triton 100X 0.2% in phosphate-buffered saline (PBS) 1X) for 30 minutes and then incubated with the corresponding antibody or lectin in blocking solution (NDS 5% and Triton 100X 0.1% in PBS 1X) overnight on an orbital shaker at 4°C. After several PBS 1X washes, sections were incubated in the corresponding secondary antibody for 2 hours at room temperature, then washed again with PBS 1X. The primary antibody used was directed against cholera toxin, B-subunit (CTB, 1/1000, goat polyclonal, Calbiochem #227040, Millipore-Merck, Guyancourt, France). The secondary antibody anti-goat was linked to the fluorochromes Alexa Fluor 594 (1/2000, Molecular Probes #A11058, Illkirch, France). For CSPGs staining, biotinylated wisteria floribunda lectin (WFA, 1/2000, Vector laboratories #B-1355, Les Ulis, France) with Alexa Fluor 488 Avidin (1/1000, Molecular Probes #A21370, Illkirch, France) was used. Images of the different sections were captured with a confocal microscope Leica TCS SPE (Nanterre Cedex, 92752 France). Images were analyzed using ImageJ 1.53n software (NIH, USA).

### Data processing and statistical analyses

The amplitude of at least 10 double integrated diaphragm EMG inspiratory bursts during normoxia and mild asphyxia was calculated for each animal from the injured and the intact sides with LabChart 8 Pro software (AD Instruments, Oxford, UNITED KINGDOM). For diaphragm EMG recordings, data obtained for right and left sides of the diaphragm of sham animals were pooled since they were not injured, and no differences were found in EMG amplitude between the two sides (data not shown, paired t-test p > 0.05).

A Student’s t-test was performed between saline and 2-DG treated groups for the extent of injury evaluation, as well as to compare diaphragm EMG amplitudes between the sham group and the intact side of C2HS animals, and between the sham group and the injured side of C2HS animals. Comparisons between the intact and injured sides for diaphragmatic EMG under normoxia, and between eupnea and asphyxia were done by Student’s paired t-test. Comparisons between the different groups for inspiratory time, expiratory time, tidal volume, respiratory rate, minute ventilation, mean arterial pressure, heart rate, surface occupied by CSPGs and CSPGs expression in the spinal cord ventral horn was evaluated by Two-way analysis of variance.

All the data were presented as mean ± SD, and statistics were considered significant when p < 0.05. SigmaPlot 12.5 software was used for all statistical analyses.

## Results

### 2-DG administration modulates ventilatory behavior

Following cervical SCI, descending respiratory pathways are impacted. Ventilation was evaluated by non-invasive plethysmography to assess the potential effects of 2-DG administration on ventilatory function recovery following cervical SCI (Figure 2A and 2B). For animals injected with saline, no difference was observed between 7d sham and 7d P.I. for all parameters: inspiratory time (228 ± 50 ms *vs* 209 ± 37 ms; p > 0.05; Figure 2C), expiratory time (404 ± 129 ms *vs* 308 ± 57 ms; p > 0.05; Figure 2D), tidal volume (0.35 ± 0.02 mL/100g *vs* 0.34 ± 0.03 mL/100g; p > 0.05; Figure 2E), respiratory rate (104 ± 23 BPM *vs* 125 ± 22 ms; p > 0.05; Figure 2F) and minute ventilation (36.01 ± 6.56 mL/min/100g *vs* 42.13 ± 7.25 mL/min/100g; p > 0.05; Figure 2G). 2-DG administration led to a significant increase in tidal volume in injured animals compared to the ones injected with saline only (0.40 ± 0.07 mL/100g *vs* 0.34 ± 0.03 mL/100g; p = 0.033). Surprisingly, sham animals treated with 2-DG also present a significant increase in tidal volume (0.46 ± 0.04 mL/100g) and minute ventilation (49.85 ± 7.49 mL/min/100g) compared to 7d sham injected with saline (p = 0.005 and p = 0.017 respectively; Figure 2E and 2G).

**Figure 2.**
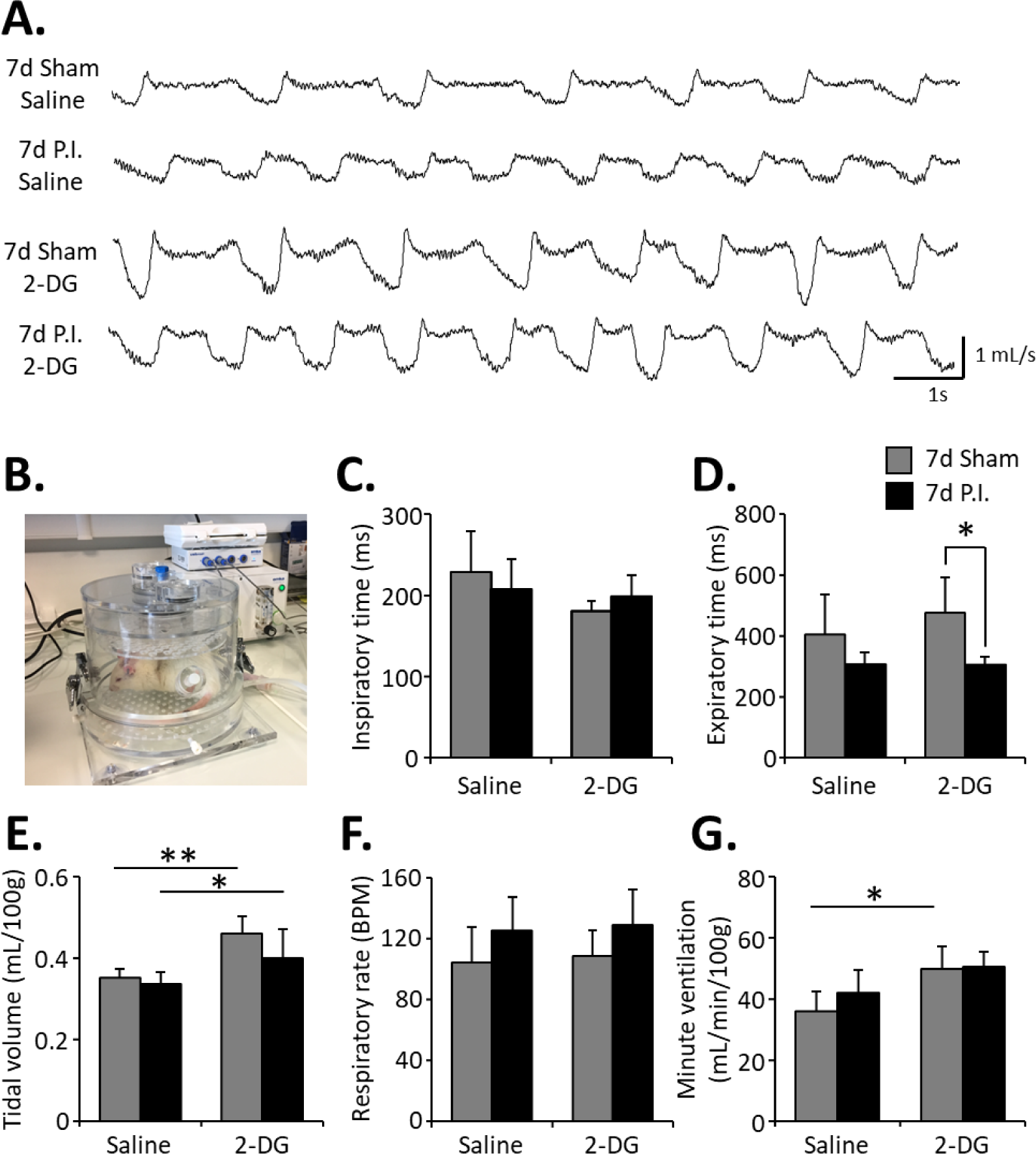
Ventilatory effects of C2HS and 2-DG administration. (A) Representative plethysmography traces at 7 days (7d) post-surgery for each group: 7d sham treated with saline or 2-deoxy-D-glucose (2-DG), and 7d post-injury (P.I.) treated with saline or 2-DG. (B) Representative picture of a rat in the plethysmography chamber (2 mL/s rate). Quantification for the different groups for (C) inspiratory time, (D) expiratory time, (E) tidal volume, (F) respiratory rate, and (G) minute ventilation. Two-way analysis of variance with Fisher LSD post-hoc analysis, * p < 0.05; ** p < 0.005. Results for tidal volume and minute ventilation have been displayed per 100g of rat. BPM = breath per minute.

### 2-DG administration has no effect on diaphragm activity

The effect of 2-DG treatment on diaphragm activity was then evaluated, (Figure 3A). During eupnea, for saline and 2-DG treated injured animals, diaphragm activity is significantly reduced on the injured side (saline group: 0.06 ± 0.07 µV.s.s.; 2-DG group: 0.05 ± 0.04 µV.s.s.) compared to the intact side (saline group: 0.54 ± 0.20 µV.s.s., p < 0.001; 2-DG group: 0.59 ± 0.13 µV.s.s., p = 0.004) and compared to sham animals (saline group: 0.37 ± 0.14 µV.s.s., p < 0.001; 2-DG group: 0.39 ± 0.19 µV.s.s., p = 0.009; Figure 3B). No effect of 2-DG was observed on diaphragm activity for all groups (p > 0.05). To study spontaneous recovery, animals were challenged with mild asphyxia (Figure 3C). For sham animals, a significant reduction in diaphragm activity during mild asphyxia compared to eupnea was observed only for saline injected group (eupnea: 0.37 ± 0.14 µV.s.s. *vs* mild asphyxia: 0.28 ± 0.13 µV.s.s.; p < 0.001). On the contrary, a significant increase during mild asphyxia was observed for both 7d P.I. groups (saline group: 0.12 ± 0.10 µV.s.s.; 2-DG group: 0.11 ± 0.04 µV.s.s.) compared to eupnea (saline group: 0.06 ± 0.07 µV.s.s., p = 0.014; 2-DG group: 0.05 ± 0.04 µV.s.s., p = 0.02).

**Figure 3.**
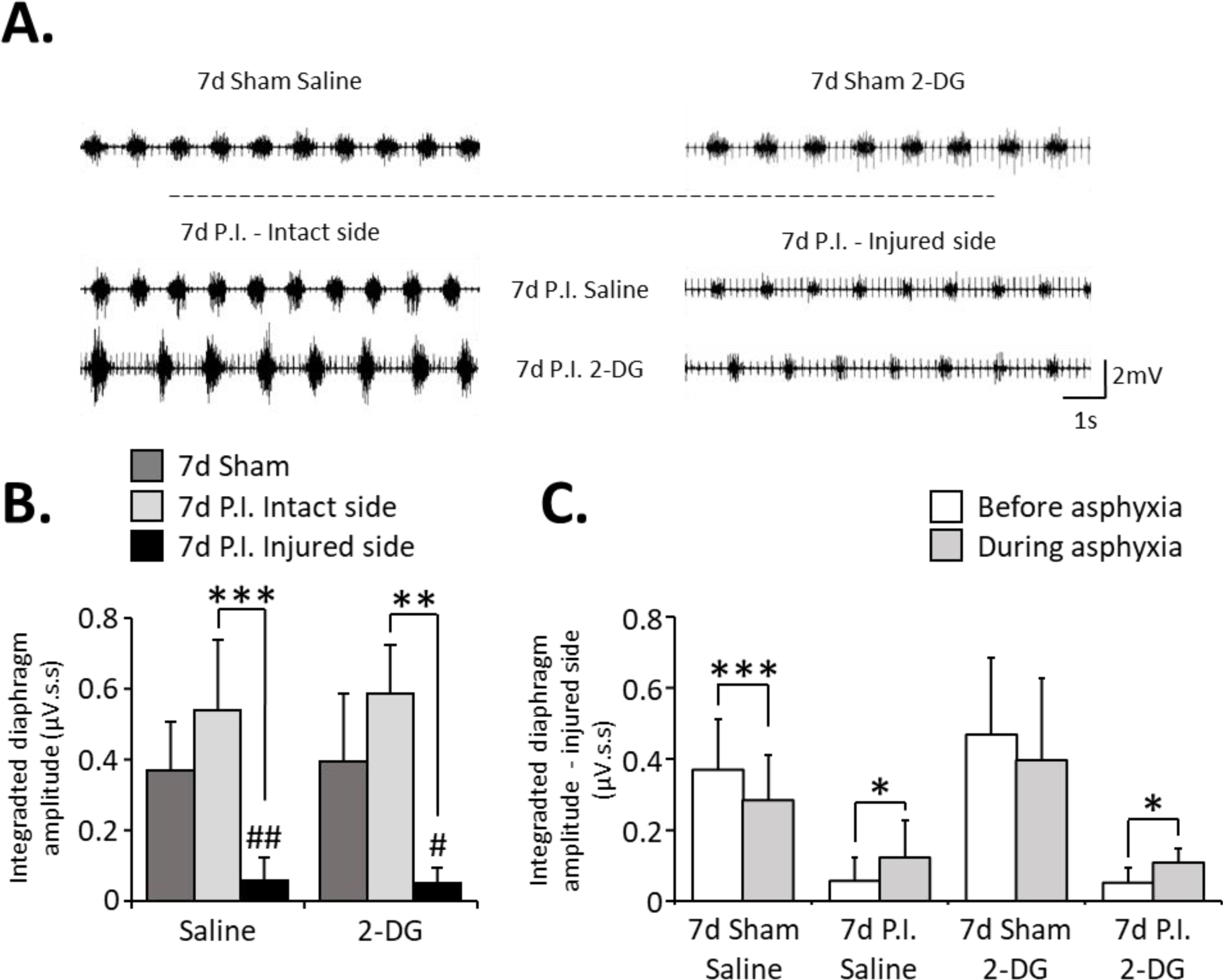
Effect of C2HS and 2-DG treatment on diaphragm activity during eupnea and mild asphyxic respiratory challenge. (A) Representative diaphragm electromyographic (EMGdia) traces for 7 days (7d) sham animals treated with saline or 2-DG, and for 7d post-injury (P.I.) groups treated with saline or 2-DG for the intact and the injured sides. (B) EMGdia amplitude quantification during eupnea. (C) EMGdia amplitude quantification during mild asphyxic respiratory challenge compared to eupnea for the sham groups and the injured side only for 7d P.I. groups. Paired t-test: * p < 0.05; ** p < 0.005; *** p < 0.001. Student’s t-test: # p < 0.005 compared to 7d sham 2-DG; ## p < 0.001 compared to 7d Sham Saline.

### Effect of 2-DG on cardiovascular function

Given the unexpected effects of 2-DG on ventilatory behavior, especially for sham groups, we wondered if 2-DG administration could have some effect on the cardiovascular function by recording femoral arterial pressure (Figure 4A). No significant difference was observed between the different groups for mean arterial pressure (MAP) (Figure 4B) and heart rate (Figure 4C) measurements, although slight increases in MAP and heart rate can be observed for the 2-DG treated sham group compared to the saline injected group (Figure 4B and 4C).

**Figure 4.**
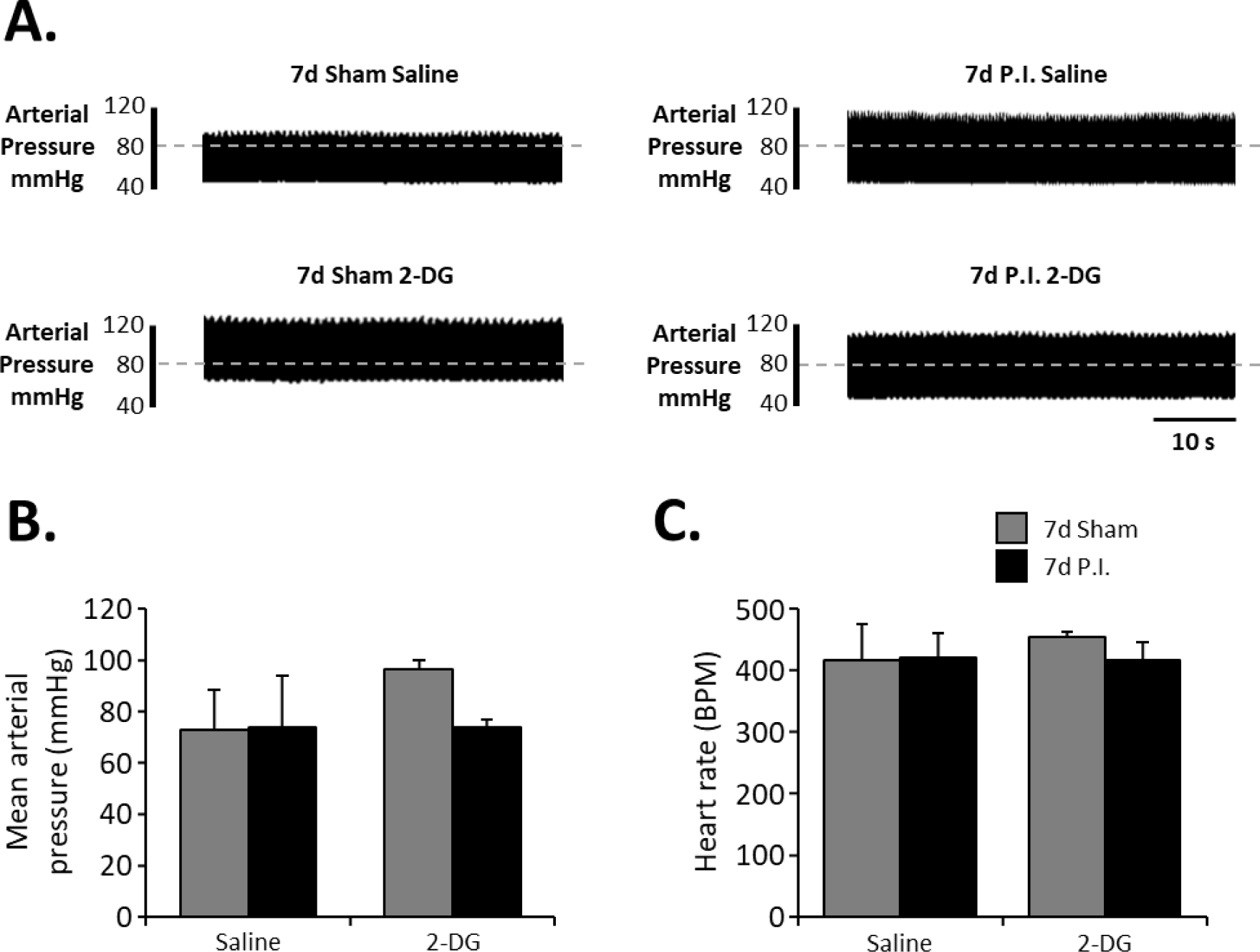
Cardiovascular effects of C2HS and 2-DG treatment at 7d post-surgery. (A) Representative traces of arterial pressure for the different groups 7 days (7d) post-surgery. (B) Mean arterial pressure (MAP) quantification for 7d sham treated with saline or 2-deoxy-D-glucose (2-DG), and 7d post-injury (P.I.) treated with saline or 2-DG. (C) Heart rate (HR) quantification for 7d sham treated with saline or (2-DG), and 7d post-injury (P.I.) treated with saline or 2-DG. Two-way analysis of variance (p > 0.05) for MAP and HR.

### Modulation of CSPGs production by 2-DG administration

Finally, we wondered if 2-DG administration could exert beneficial effects on inflammatory processes in the spinal cord following SCI, particularly those involved in inhibiting neuronal plasticity, such as the production of CSGPs from the perineuronal net. We evaluated CSPGs production specifically around CTB-labeled phrenic motoneurons in the C3-C6 spinal cord ventral horn (Figure 5A and B). A C2HS leads to a significant increase in the surface area occupied by CSPGs compared to sham animals when injected with saline (17.63 ± 5.17 % *vs* 4.71 ± 0.62 % respectively; p < 0.001). Among 7d P.I. animals, the 2-DG treatment does not lead to any change in CSPGs production. However, 7d sham animals treated with 2-DG demonstrate a significant increase in surface occupied by CSPGs labeling (4.71 ± 0.62 % *vs* 28.37 ± 4.50 % respectively; p = 0.08; Figure 5C) and CSPGs intensity (114.74 ± 1.49 AU *vs* 137.03 ± 3.66 AU respectively; p = 0.011; Figure 5D). Interestingly, 2-DG treated animals that received a C2HS versus the sham surgery present a significantly lower surface occupied by CSPGs expression (13.68 ± 4.08 % *vs* 28.37 ± 4.50 % respectively; p = 0.02; Figure 5C) and CSPGs intensity (118.30 ± 5.08 AU *vs* 137.03 ± 3.66 AU respectively; p = 0.011; Figure 5D).

**Figure 5.**
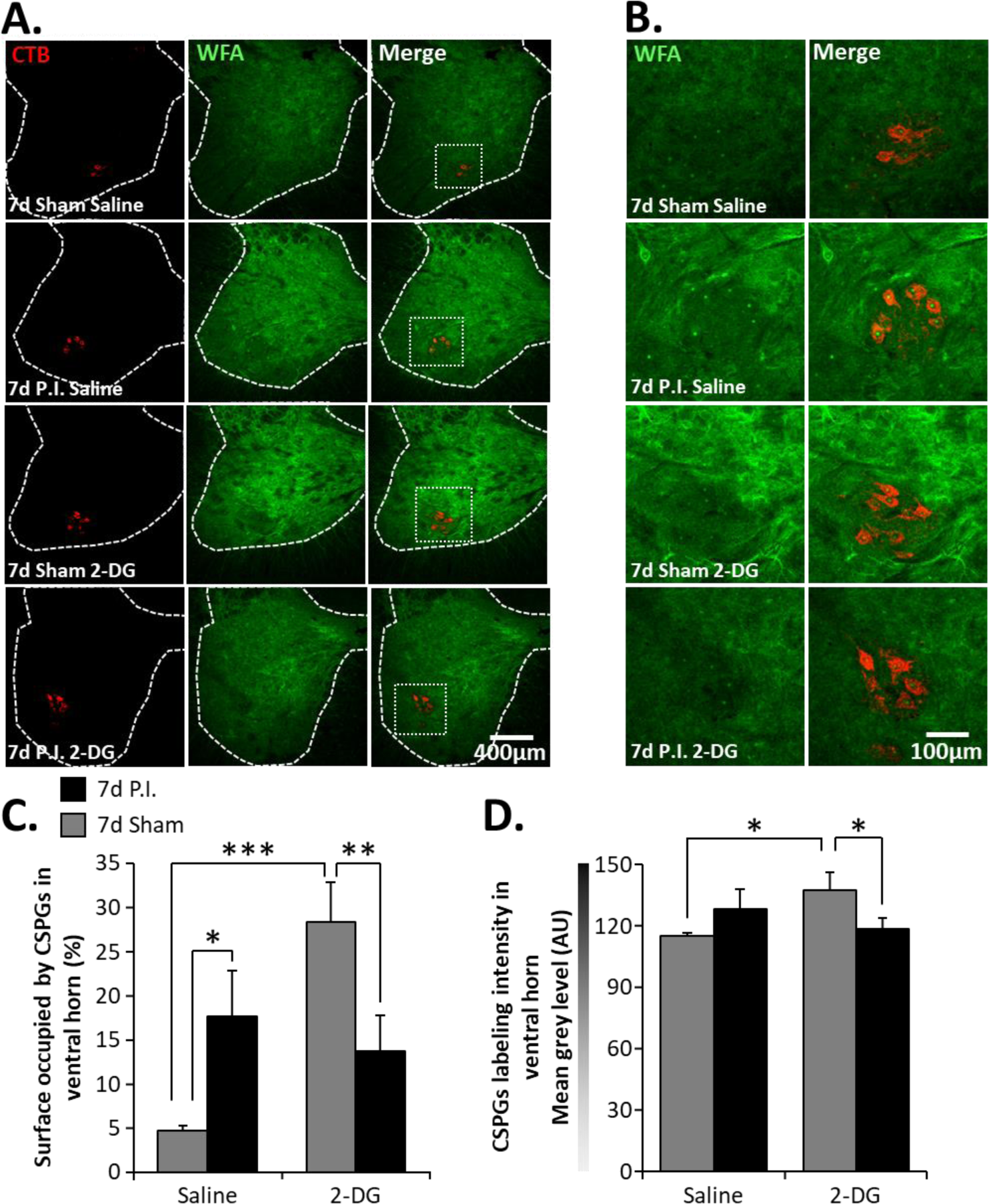
Effect of C2HS and 2-DG administration on CSPGs production. (A) Representative pictures showing the location of phrenic motoneurons in red and CSPGs (WFA labeling) in green in the C3-C6 spinal cord ventral horn at 7 days (7d) sham treated with saline or 2-deoxy-D-glucose (2-DG), and at 7d post-injury (P.I.) treated with saline or 2-DG. (B) Magnification of phrenic motoneurons surrounded by CSPGs displayed in A. Note the presence of CSPGs around phrenic motor pools, especially for 7d Sham 2-DG group. (C) Quantification of the surface occupied by CSPGs in the spinal cord ventral horn for the different groups at 7d post-surgery. (D) CSPGs intensity quantification in the spinal cord ventral horn for each group at 7d post-surgery. Expression was determined by quantifying the grey level corresponding to WFA labeling intensity. Two-way analysis of variance with Fisher LSD post-hoc analysis, * p < 0.05; ** p < 0.005; *** p < 0.001.

## Discussion

The present study is the first to explore the potential therapeutic effects of 2-DG for enhancing respiratory function after cervical spinal cord injury (SCI) in a preclinical model. A previous study employed 2-DG as a glycolysis inhibitor following SCI, but only to demonstrate that rostro-caudal disparities in energy metabolism leads to differences in rostro-caudal axonal degeneration spreading in early thoracic (T10) SCI in mice (Ohnishi et al., 2021). This work showcased the importance of metabolism (glycolysis in particular) in post-injury events occurring in the spinal cord, supporting our hypothesis that 2-DG treatment could lead to beneficial effects on functional recovery following SCI.

At 7d P.I., injured animals injected with saline showed no significant changes compared to sham animals also injected with saline in terms of ventilatory behavior and cardiovascular function. It was previously reported that MAP is initially reduced following injury (Vinit et al., 2016; Michel-Flutot et al., 2023b), but returns to pre-injury values by 7d P.I. (Vinit et al., 2016; Lee and Gonzalez-Rothi, 2017; Michel-Flutot et al., 2023b), despite the persistence of systolic dysfunction (Michel-Flutot et al., 2023b). This observation could be due to vascular compensations occurring between the moment of injury and 7d P.I., such as reorganization of the sympathetic neuronal network (i.e., around preganglionic neurons) (Krassioukov and Weaver, 1996). Concerning ventilatory behavior, no dysfunction was observed in 7d injured animals when evaluated under normal breathing using plethysmography as previously shown (Michel-Flutot et al., 2023b). However, other studies have demonstrated initial modifications occur in some respiratory parameters following injury, including a decrease in tidal volume and minute ventilation, associated with an augmentation in respiratory frequency (Golder et al., 2001b; Lovett-Barr et al., 2012; Navarrete-Opazo et al., 2017). In this model, a spontaneous return to pre-injury values over time has been observed (Navarrete-Opazo et al., 2017). This spontaneous recovery can be explained by the recruitment of additional respiratory-related muscles to compensate for the loss of hemidiaphragm activity on the injured side in this C2HS preclinical model (Katagiri et al., 1994; Fuller et al., 2008; Keomani et al., 2014; Michel-Flutot et al., 2022). This effect allows the animals to maintain their respiratory gas exchange homeostasis and keeps respiratory function similar to that of non-injured animals (Beth Zimmer et al., 2015; Navarrete-Opazo et al., 2015). As expected, the C2HS also led to an upregulation of CSPGs in the spinal cord tissue (Lemons et al., 1999), including the area immediately surrounding phrenic motoneurons. The CSPGs upregulation could in part explain why no recovery of spontaneous diaphragm activity can be achieved physiologically and spontaneously after high SCI, especially when considering that removal of CSPGs leads to a recovery of respiratory function in a C2HS model (Warren et al., 2018).

Interestingly, contrary to our initial hypothesis, 2-DG treatment in injured animals did not induce any beneficial effects regarding diaphragm activity (during eupnea and asphyxic challenge), or CSPGs production compared to injured animals injected with saline. This, however, surprisingly led to a significant increase in tidal volume and a subsequent increase in minute ventilation for these 2-DG treated animals. Unexpectedly, sham animals injected with the same dose of 2-DG presented with the similar increases in tidal volume and minute ventilation. During asphyxia, 2-DG treated sham do not present a significant decrease in diaphragm EMG amplitude, which is likely attributable to a wider variability in animal response to asphyxia compared to saline injected sham animals. This could be related to global modifications in breathing behavior when sham animals are treated with 2-DG. Furthermore, a significant increase in the surface occupied by CSPGs and CSPGs intensity in the C3-C6 spinal cord ventral horn can be observed compared to sham injected with saline. This phenomenon could result from inflammatory processes in the spinal cord following 2-DG treatment, which may involve the activation of astrocytes and successive overproduction of CSPGs. For example, *in vitro* studies have shown that oxygen-glucose deprivation leads to astrocyte activation, and subsequent CSPGs overproduction (Li et al., 2021). An inflammatory state due to 2-DG could also explain the increased ventilation observed in these animals. Inflammatory processes are known to increase oxygen demands due to the energetic requirements of cells involved in inflammation and immunological response (Zenewicz, 2017). Inflammatory signals also induce plasticity in central nervous system respiratory centers, impacting the activity of respiratory-related neurons, which can then modify ventilatory behavior (Hocker et al., 2017). Here, given the absence of change in diaphragm EMG amplitude, we can hypothesize that the increase in tidal volume and minute ventilation in sham rats treated with 2-DG results from the recruitment of respiratory accessory muscles to handle the increase in oxygen needs due to inflammation. This could be further explored using metabolic plethysmography to evaluate the levels of oxygen consumed and CO_2_ produced.

These results remain surprising considering that we used a daily dose (200 mg/kg/day) lower than ones used in several previous studies. Interestingly, higher doses of 2-DG (500 mg/kg/day) do not have adverse effects on learning and memory functions in adult rats (Ockuly et al., 2012). 2-DG has also showed beneficial effects in preclinical studies as an antiepileptic and anticonvulsant without inducing observable deleterious effects when delivered at doses ranging from 37.5 mg/kg to 500 mg/kg over a prolonged period of time (Stafstrom et al., 2008; Stafstrom et al., 2009; Rho et al., 2019; Sutula et al., 2023). However, it has been shown that 2-DG could lead to cardiac toxicity, associated with body weight loss and reduced survival rate when administered chronically (Minor et al., 2010). This could explain the increased MAP values we observed in sham animals treated with 2-DG. This however remains to be confirmed in our treatment paradigm for the dose we administered to our animals.

The different responses to 2-DG we observed between C2HS and sham animals could be explained by the difference in inflammatory state between the groups. C2HS animals are already undergoing active inflammation induced by SCI, while the sham animals underwent the same surgery without damaging the spinal cord. We can hypothesize that 2-DG induces a new inflammatory state in sham animals by modifying their metabolic physiology. This does not happen in C2HS animals since an inflammatory state is already present. This could mean that 2-DG could still be of use following SCI. The lack of effect we observe in C2HS animals in our study could stem from an inadequate dose of 2-DG, a treatment duration that was too short, or a combination of both. A future study could answer these questions by testing several doses of 2-DG with longer treatments intervals following SCI in order to optimize the potential effect. Further supporting this, administration of 2-DG has also been used to demonstrate the involvement of glycolytic flux in neurorespiratory plasticity induced by intermittent hypoxia. More precisely, phrenic long-term facilitation induced by moderate acute intermittent hypoxia in healthy rats preconditioned with repetitive acute intermittent hypoxia is abolished in animals that received 2-DG orally (MacFarlane et al., 2017). This highlights the importance of neuronal glycolytic activity for respiratory plasticity in intact/healthy animals.

In conclusion, this study is the first to evaluate the potential beneficial effects of 2-DG on neuroinflammation and respiratory function following high SCI. While 2-DG treatment had no effect on injured animals at the dose we used, it had surprising deleterious physiological and inflammatory effects in non-injured animals. The absence of deleterious effect in injured animals lead us to believe that 2-DG could still be a potential therapeutic for SCI repair. Future studies using higher doses of 2-DG administered over a longer period of time could unravel potential beneficial effects of 2-DG on neuroinflammatory processes occurring after SCI.

## Authors Contribution

SV designed research; PMF and SV performed research; PMF analyzed data; PMF, AM and SVI wrote the paper. All authors have read and agreed to the published version of the manuscript.

## Acknowledgements

The authors thank Samantha (Sammi) J. Thomas for editing the present manuscript. In This study has benefited from the facilities of CYMAGES and histology (UFR SVS, UVSQ, Université Paris-Saclay, 78180 Montigny-le-Bretonneux, France).

## References

Ahuja CS, Wilson JR, Nori S, Kotter MRN, Druschel C, Curt A, Fehlings MG (2017) Traumatic spinal cord injury. Nature Reviews Disease Primers 3:17018.

Allen LL, Nichols NL, Asa ZA, Emery AT, Ciesla MC, Santiago JV, Holland AE, Mitchell GS, Gonzalez-Rothi EJ (2021) Phrenic motor neuron survival below cervical spinal cord hemisection. Experimental neurology 346:113832.

Bartus K, James ND, Bosch KD, Bradbury EJ (2012) Chondroitin sulphate proteoglycans: key modulators of spinal cord and brain plasticity. Experimental neurology 235:5–17.

Beth Zimmer M, Grant JS, Ayar AE, Goshgarian HG (2015) Ipsilateral inspiratory intercostal muscle activity after C2 spinal cord hemisection in rats. J Spinal Cord Med 38:224–230.

Bradbury EJ, Burnside ER (2019) Moving beyond the glial scar for spinal cord repair. Nat Commun 10:3879.

Bradbury EJ, Moon LD, Popat RJ, King VR, Bennett GS, Patel PN, Fawcett JW, McMahon SB (2002) Chondroitinase ABC promotes functional recovery after spinal cord injury. Nature 416:636–640.

Cregg JM, DePaul MA, Filous AR, Lang BT, Tran A, Silver J (2014) Functional regeneration beyond the glial scar. Experimental neurology 253:197–207.

Dey S, Murmu N, Mondal T, Saha I, Chatterjee S, Manna R, Haldar S, Dash SK, Sarkar TR, Giri B (2022) Multifaceted entrancing role of glucose and its analogue, 2-deoxy-D-glucose in cancer cell proliferation, inflammation, and virus infection. Biomedicine & Pharmacotherapy 156:113801.

Dickendesher TL, Baldwin KT, Mironova YA, Koriyama Y, Raiker SJ, Askew KL, Wood A, Geoffroy CG, Zheng B, Liepmann CD, Katagiri Y, Benowitz LI, Geller HM, Giger RJ (2012) NgR1 and NgR3 are receptors for chondroitin sulfate proteoglycans. Nature neuroscience 15:703–712.

Dougherty BJ, Lee KZ, Lane MA, Reier PJ, Fuller DD (2012) Contribution of the spontaneous crossed-phrenic phenomenon to inspiratory tidal volume in spontaneously breathing rats. J Appl Physiol (1985) 112:96-105.

Fawcett JW, Schwab ME, Montani L, Brazda N, Müller HW (2012) Defeating inhibition of regeneration by scar and myelin components. Handb Clin Neurol 109:503–522.

Fayssoil A, Michel-Flutot P, Lofaso F, Carlier R, El Hajjam M, Vinit S, Mansart A (2021) Analysis of inspiratory and expiratory muscles using ultrasound in rats: A reproducible and non-invasive tool to study respiratory function. Respir Physiol Neurobiol 285:103596.

Fisher D, Xing B, Dill J, Li H, Hoang HH, Zhao Z, Yang X-L, Bachoo R, Cannon S, Longo FM, Sheng M, Silver J, Li S (2011) Leukocyte Common Antigen-Related Phosphatase Is a Functional Receptor for Chondroitin Sulfate Proteoglycan Axon Growth Inhibitors. The Journal of Neuroscience 31:14051–14066.

Fuller DD, Golder FJ, Olson EB, Jr., Mitchell GS (2006) Recovery of phrenic activity and ventilation after cervical spinal hemisection in rats. J Appl Physiol (1985) 100:800-806.

Fuller DD, Doperalski NJ, Dougherty BJ, Sandhu MS, Bolser DC, Reier PJ (2008) Modest spontaneous recovery of ventilation following chronic high cervical hemisection in rats. Exp Neurol 211:97–106.

Ghali MGZ (2017) The crossed phrenic phenomenon. Neural Regen Res 12:845–864.

Golder FJ, Reier PJ, Bolser DC (2001a) Altered respiratory motor drive after spinal cord injury: Supraspinal and bilateral effects of a unilateral lesion. J Neurosci 21:8680–8689.

Golder FJ, Reier PJ, Davenport PW, Bolser DC (2001b) Cervical spinal cord injury alters the pattern of breathing in anesthetized rats. J Appl Physiol 91:2451–2458.

Goshgarian HG (2009) The crossed phrenic phenomenon and recovery of function following spinal cord injury. Respiratory physiology & neurobiology 169:85–93.

Hocker AD, Stokes JA, Powell FL, Huxtable AG (2017) The impact of inflammation on respiratory plasticity. Experimental neurology 287:243–253.

Ip WKE, Hoshi N, Shouval DS, Snapper S, Medzhitov R (2017) Anti-inflammatory effect of IL-10 mediated by metabolic reprogramming of macrophages. Science 356:513–519.

James ND, Shea J, Muir EM, Verhaagen J, Schneider BL, Bradbury EJ (2015) Chondroitinase gene therapy improves upper limb function following cervical contusion injury. Experimental neurology 271:131–135.

Katagiri M, Young RN, Platt RS, Kieser TM, Easton PA (1994) Respiratory muscle compensation for unilateral or bilateral hemidiaphragm paralysis in awake canines. J Appl Physiol 77:1972–1982.

Keomani E, Deramaudt TB, Petitjean M, Bonay M, Lofaso F, Vinit S (2014) A murine model of cervical spinal cord injury to study post-lesional respiratory neuroplasticity. J Vis Exp 87:e51235.

Krassioukov AV, Weaver LC (1996) Morphological changes in sympathetic preganglionic neurons after spinal cord injury in rats. Neuroscience 70:211–225.

Kwok JCF, Heller JP, Zhao R-R, Fawcett JW (2014) Targeting Inhibitory Chondroitin Sulphate Proteoglycans to Promote Plasticity After Injury. In: Axon Growth and Regeneration: Methods and Protocols (Murray AJ, ed), pp 127–138. New York, NY: Springer New York.

Lane MA, Lee K-Z, Fuller DD, Reier PJ (2009) Spinal circuitry and respiratory recovery following spinal cord injury. Respiratory physiology & neurobiology 169:123–132.

Lee K-Z, Gonzalez-Rothi EJ (2017) Contribution of 5-HT2A receptors on diaphragmatic recovery after chronic cervical spinal cord injury. Respiratory physiology & neurobiology 244:51–55.

Lee KZ, Huang YJ, Tsai IL (2014) Respiratory motor outputs following unilateral midcervical spinal cord injury in the adult rat. J Appl Physiol 116:395–405.

Lemons ML, Howland DR, Anderson DK (1999) Chondroitin Sulfate Proteoglycan Immunoreactivity Increases Following Spinal Cord Injury and Transplantation. Experimental neurology 160:51–65.

Li S, Duan Q, Lu M, Wen X, Chen J, Tan S, Guo Y (2021) CSPGs promote the migration of meningeal fibroblasts via p38 mitogen-activated protein kinase signaling pathway under OGD conditions. Brain Research Bulletin 173:37–44.

Locke KC, Randelman ML, Hoh DJ, Zholudeva LV, Lane MA (2022) Respiratory plasticity following spinal cord injury: perspectives from mouse to man. Neural Regen Res 17:2141–2148.

Lovett-Barr MR, Satriotomo I, Muir GD, Wilkerson JE, Hoffman MS, Vinit S, Mitchell GS (2012) Repetitive intermittent hypoxia induces respiratory and somatic motor recovery after chronic cervical spinal injury. J Neurosci 32:3591–3600.

MacFarlane PM, Vinit S, Mitchell GS (2017) Enhancement of phrenic long-term facilitation following repetitive acute intermittent hypoxia is blocked by the glycolytic inhibitor 2-deoxyglucose. Am J Physiol Regul Integr Comp Physiol:ajpregu 00306 02017.

Mantilla CB, Zhan WZ, Sieck GC (2009) Retrograde labeling of phrenic motoneurons by intrapleural injection. J Neurosci Methods 182:244–249.

Michel-Flutot P, Lane MA, Lepore AC, Vinit S (2023a) Therapeutic Strategies Targeting Respiratory Recovery after Spinal Cord Injury: From Preclinical Development to Clinical Translation. Cells 12:1519.

Michel-Flutot P, Mansart A, Fayssoil A, Vinit S (2023b) Effects of C2 hemisection on respiratory and cardiovascular functions in rats. Neural Regen Res 18.

Michel-Flutot P, Jesus I, Vanhee V, Bourcier CH, Emam L, Ouguerroudj A, Lee K-Z, Zholudeva LV, Lane MA, Mansart A, Bonay M, Vinit S (2022) Effects of Chronic High-Frequency rTMS Protocol on Respiratory Neuroplasticity Following C2 Spinal Cord Hemisection in Rats. Biology 11:473.

Minor RK, Smith DL, Jr., Sossong AM, Kaushik S, Poosala S, Spangler EL, Roth GS, Lane M, Allison DB, de Cabo R, Ingram DK, Mattison JA (2010) Chronic ingestion of 2-deoxy-D-glucose induces cardiac vacuolization and increases mortality in rats. Toxicol Appl Pharmacol 243:332–339.

Muir E, De Winter F, Verhaagen J, Fawcett J (2019) Recent advances in the therapeutic uses of chondroitinase ABC. Experimental neurology 321:113032.

Nantwi KD, El-Bohy AA, Schrimsher GW, Reier PJ, Goshgarian HG (1999) Spontaneous Functional Recovery in a Paralyzed Hemidiaphragm Following Upper Cervical Spinal Cord Injury in Adult Rats. Neurorehabilitation and Neural Repair 13:225–234.

Navarrete-Opazo A, Dougherty BJ, Mitchell GS (2017) Enhanced recovery of breathing capacity from combined adenosine 2A receptor inhibition and daily acute intermittent hypoxia after chronic cervical spinal injury. Exp Neurol 287:93–101.

Navarrete-Opazo A, Vinit S, Dougherty BJ, Mitchell GS (2015) Daily acute intermittent hypoxia elicits functional recovery of diaphragm and inspiratory intercostal muscle activity after acute cervical spinal injury. Exp Neurol 266:1–10.

Obaid M, Udden SMN, Alluri P, Mandal SS (2021) LncRNA HOTAIR regulates glucose transporter Glut1 expression and glucose uptake in macrophages during inflammation. Scientific reports 11:232.

Ockuly JC, Gielissen JM, Levenick CV, Zeal C, Groble K, Munsey K, Sutula TP, Stafstrom CE (2012) Behavioral, cognitive, and safety profile of 2-deoxy-2-glucose (2DG) in adult rats. Epilepsy Res 101:246–252.

Ohnishi Y, Yamamoto M, Sugiura Y, Setoyama D, Kishima H (2021) Rostro-caudal different energy metabolism leading to differences in degeneration in spinal cord injury. Brain Commun 3:fcab058.

Pajak B, Siwiak E, Sołtyka M, Priebe A, Zieliński R, Fokt I, Ziemniak M, Jaśkiewicz A, Borowski R, Domoradzki T, Priebe W (2020) 2-Deoxy-d-Glucose and Its Analogs: From Diagnostic to Therapeutic Agents. International journal of molecular sciences 21:234.

Porter WT (1895) The Path of the Respiratory Impulse from the Bulb to the Phrenic Nuclei. J Physiol 17:455–485.

Rana S, Zhan W-Z, Sieck GC, Mantilla CB (2022) Cervical spinal hemisection alters phrenic motor neuron glutamatergic mRNA receptor expression. Experimental neurology:114030.

Randelman M, Zholudeva LV, Vinit S, Lane MA (2021) Respiratory Training and Plasticity After Cervical Spinal Cord Injury. Frontiers in Cellular Neuroscience 15.

Rho JM, Shao LR, Stafstrom CE (2019) 2-Deoxyglucose and Beta-Hydroxybutyrate: Metabolic Agents for Seizure Control. Front Cell Neurosci 13:172.

Rosenzweig ES et al. (2019) Chondroitinase improves anatomical and functional outcomes after primate spinal cord injury. Nature neuroscience 22:1269–1275.

Shen Y, Tenney AP, Busch SA, Horn KP, Cuascut FX, Liu K, He Z, Silver J, Flanagan JG (2009) PTPsigma is a receptor for chondroitin sulfate proteoglycan, an inhibitor of neural regeneration. Science 326:592–596.

Stafstrom CE, Roopra A, Sutula TP (2008) Seizure suppression via glycolysis inhibition with 2-deoxy-D-glucose (2DG). Epilepsia 49 Suppl 8:97–100.

Stafstrom CE, Ockuly JC, Murphree L, Valley MT, Roopra A, Sutula TP (2009) Anticonvulsant and antiepileptic actions of 2-deoxy-D-glucose in epilepsy models. Ann Neurol 65:435–447.

Sutula TP, Wilson ST, Franzoso S, Stafstrom CE (2023) 2-Deoxy-D-glucose administration after seizures has disease-modifying effects on kindling progression. Epilepsy Res 193:107169.

Tester NJ, Howland DR (2008) Chondroitinase ABC improves basic and skilled locomotion in spinal cord injured cats. Experimental neurology 209:483–496.

Vinit S, Kastner A (2009) Descending bulbospinal pathways and recovery of respiratory motor function following spinal cord injury. Respir Physiol Neurobiol 169:115–122.

Vinit S, Gauthier P, Stamegna JC, Kastner A (2006) High cervical lateral spinal cord injury results in long-term ipsilateral hemidiaphragm paralysis. J Neurotrauma 23:1137–1146.

Vinit S, Keomani E, Deramaudt TB, Bonay M, Petitjean M (2016) Reorganization of Respiratory Descending Pathways following Cervical Spinal Partial Section Investigated by Transcranial Magnetic Stimulation in the Rat. PloS one 11:e0148180.

Wang Q, Zhao Y, Sun M, Liu S, Li B, Zhang L, Yang L (2014) 2-Deoxy-d-glucose attenuates sevoflurane-induced neuroinflammation through nuclear factor-kappa B pathway in vitro. Toxicology in vitro : an international journal published in association with BIBRA 28:1183–1189.

Warren PM, Steiger SC, Dick TE, MacFarlane PM, Alilain WJ, Silver J (2018) Rapid and robust restoration of breathing long after spinal cord injury. Nat Commun 9:4843–4843.

Winslow C, Rozovsky J (2003) Effect of spinal cord injury on the respiratory system. Am J Phys Med Rehabil 82:803–814.

Woodward GE, Hudson MT (1954) The effect of 2-desoxy-D-glucose on glycolysis and respiration of tumor and normal tissues. Cancer Res 14:599–605.

Xi H, Kurtoglu M, Lampidis TJ (2014) The wonders of 2-deoxy-D-glucose. IUBMB Life 66:110–121.

York EM, Zhang J, Choi HB, MacVicar BA (2021) Neuroinflammatory inhibition of synaptic long-term potentiation requires immunometabolic reprogramming of microglia. Glia 69:567–578.

Zenewicz LA (2017) Oxygen Levels and Immunological Studies. Frontiers in Immunology 8.

